# Multiple roles of the non-structural protein 3 (nsP3) alphavirus unique domain (AUD) during Chikungunya virus genome replication and transcription

**DOI:** 10.1101/377127

**Authors:** Yanni Gao, Niluka Goonawardane, Andrew Tuplin, Mark Harris

## Abstract

Chikungunya virus (CHIKV) is a re-emerging *Alphavirus* causing fever, joint pain, skin rash, arthralgia, and occasionally death. Antiviral therapies and/or effective vaccines are urgently required. CHIKV biology is poorly understood, in particular the functions of the non-structural protein 3 (nsP3). Here we present the results of a mutagenic analysis of the alphavirus unique domain (AUD) of nsP3. Informed by the structure of the Sindbis virus AUD and an alignment of amino acid sequences of multiple alphaviruses, a series of mutations in the AUD were generated in a CHIKV sub-genomic replicon. This analysis revealed an essential role for the AUD in CHIKV RNA replication, with mutants exhibiting species- and cell-type specific phenotypes. To test if the AUD played a role in other stages of the virus lifecycle, the mutant panel was also analysed in the context of infectious CHIKV. Results indicated that, in addition to a role in RNA replication, the AUD was also required for virus assembly. Further analysis revealed that one mutant (P247A/V248A) specifically blocked transcription of the subgenomic RNA leading to a dramatic reduction in synthesis of the structural proteins and concomitant reduction in virus production. This phenotype could be explained by both a reduction in the binding of the P247A/V248A mutant nsP3 to viral genomic RNA *in vivo*, and the reduced affinity of the mutant AUD for the subgenomic promoter RNA *in vitro.* We propose that the AUD is a pleiotropic protein domain, with multiple functions during CHIKV RNA synthesis.

**Author summary:** Chikungunya virus (CHIKV) is an emerging threat to world health. It is transmitted by Aedes species mosquitos, and has caused massive epidemics across the globe. The virus causes fever, rash, arthritis and can sometimes be fatal. The biology of CHIKV is poorly understood, to address this deficiency we aimed to identify functions of one of the viral proteins, nsP3. We focussed on the central part of this protein, termed the alphavirus unique domain (AUD) because it is unique to the genus of viruses to which CHIKV belongs – the *Alphaviruses* – and not present in other related viruses. By making changes (mutations) in the AUD and analysing the effects of these changes we show that it is involved in multiple stages of the virus lifecycle. These observations identify nsP3 and the AUD in particular as a potential target for antiviral therapy or rational vaccine design.

## Introduction

Chikungunya virus (CHIKV; family *Togaviridae*, genus *Alphavirus)* [1] is an arbovirus that causes fever, rash and arthralgia with an infrequent fatal outcome [2]. It was first isolated in Tanzania in 1952-1953 [3, 4]. During the last 50 years, numerous CHIKV re-emergences have been documented across the world, including Africa, Asia, Europe and America [5, 6]. CHIKV is transmitted to humans by mosquitoes, mainly *Aedes aegypti* and *Ae. albopictus.* The latter can reproduce in more moderate climates which means that CHIKV has spread from Southern Africa and is now present across the Americas and parts of Southern Europe (including France and Italy). Increasing global temperatures resulting from climate change raise the concern that CHIKV will spread further. In this regard, there are no antiviral therapies or safe, effective vaccines available to treat CHIKV infection.

CHIKV has an 11.5 kilobase positive-sense, single-stranded RNA genome that is both capped and polyadenylated, and contains two open reading frames (ORFs). The first ORF is translated directly from full-length genomic RNA and encodes the non-structural proteins nsP1 to nsP4. These four proteins are required for RNA synthesis – generating both negative and positive full-length genomic RNA and a smaller subgenomic RNA from which the second ORF is translated to yield the structural proteins (capsid, envelope glycoproteins E1-3 and the 6K viroporin). Biochemical functions have been ascribed to 3 of the nsPs: nsP1 exhibits methyl- and guanyl-transferase activities, nsP2 is a helicase/protease, and nsP4 is the RNA-dependent

RNA polymerase. Although nsP3 plays an essential role in RNA replication, its biochemical functions remain largely undefined [7, 8]. It is proposed to comprise three domains: at the N-terminus is a macro-domain which exhibits both ADP-ribose and RNA binding, and ADP-ribosylhydrolase capabilities [9–11]. This is followed by the alphavirus unique domain (AUD), so called as it is only present in the *Alphavirus* genus and is absent from the closely related Rubella virus (the sole member of the *Rubivirus* genus within the *Togaviridae)*, and a C-terminal hypervariable region (Fig. 1A). The latter plays an important role in virus-host interactions and may be a significant determinant of pathogenesis through interactions with cell-type-specific factors [12, 13]. The AUD is located in the centre of nsP3, and despite a high level of sequence homology across the alphaviruses, the function of this domain remains elusive. Of note, the structure of the Sindbis virus (SINV) AUD has been determined in the context of a pre-cleavage fragment of the polyprotein spanning the C-terminus of nsP2 (protease and methyl-transferase-like domains), and the N-terminus of nsP3 (macrodomain and AUD) [14]. This revealed that the AUD presents a unique protein fold containing a zinc coordination site. In this study we sought to investigate the function of AUD during the virus lifecycle in cells derived from both the vertebrate host and the mosquito vector, to identify targets for antiviral intervention and means of rational attenuation for vaccine development. By mutagenic analysis we demonstrate that the AUD exhibits both species- and cell-type specific phenotypes, and plays roles in both virus genome replication and structural protein expression.

**Figure 1.**
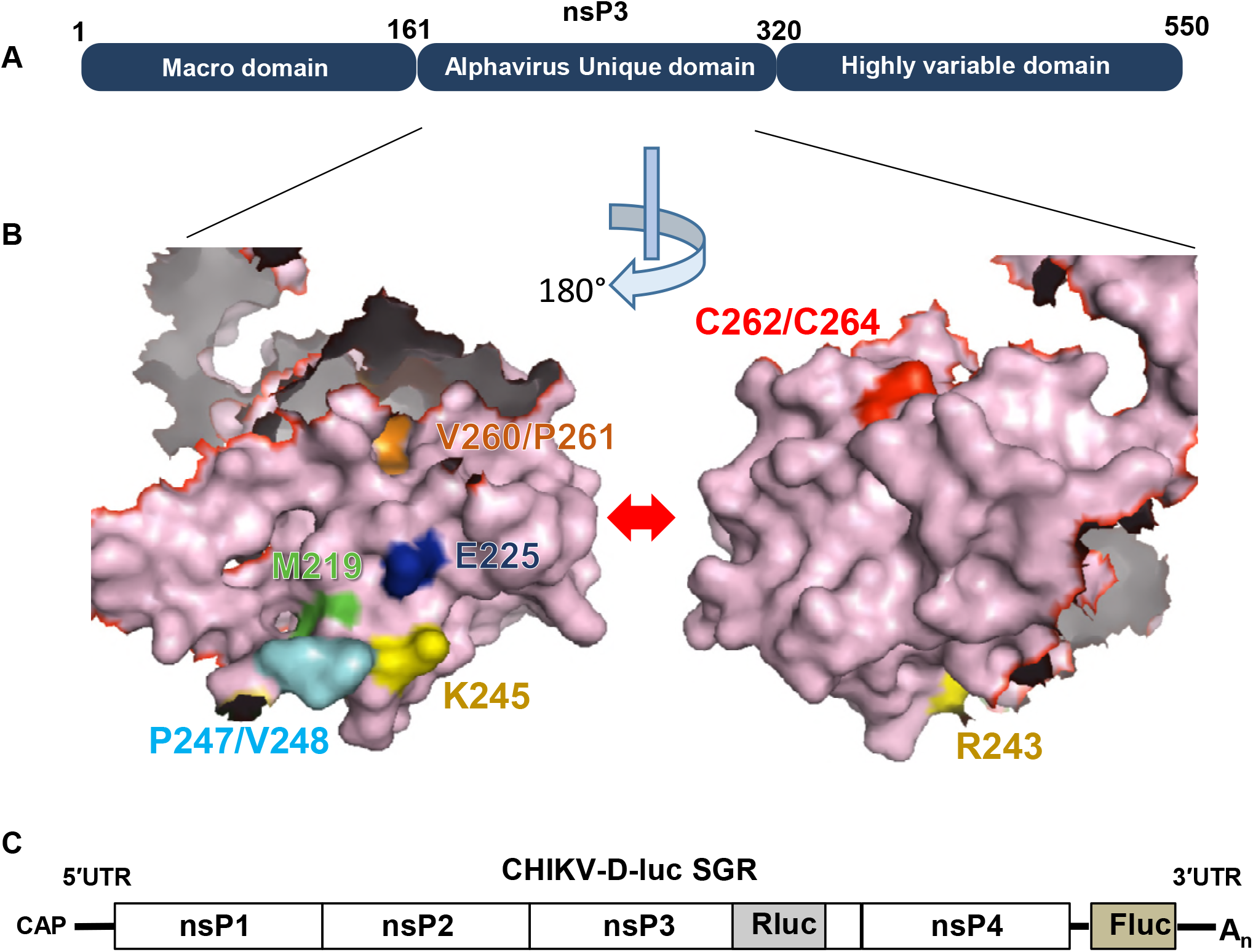
(A) Three domain structure of the alphavirus nsP3 protein. (B) Surface representation of the Sindbis virus nsP3 AUD structure (PDB ID code 4GUA) [14] (residues 161-320), including the 40 amino acid flexible linker between the macrodomain and the AUD. The locations of the mutated residues in nsP3 are indicated. The two images show opposite faces of the structure, rotated 180° along the vertical axis. (C) Structure of CHIKV-D-Luc-SGR. (RLuc: Renilla luciferase, FLuc: firefly luciferase) RLuc is expressed as an internal fusion with nsP3 and thus is produced following translation of the input RNA. RLuc activity is therefore an indirect measure of both input translation and replication. FLuc is expressed from the subgenomic promoter and thus is only produced after RNA replication has occurred.

## Results

### Construction of CHIKV subgenomic replicons with AUD mutations

To identify residues within the AUD that are conserved across the *Alphavirus* genus we first aligned the AUD amino acid sequences of a range of both Old World and New World alphaviruses (Supplementary Fig S1A). As the AUD sequences between SINV and CHIKV are highly conserved (118 of 243 residues are identical), the nsP2/nsP3 protein structure of SINV [14] was referenced to identify the putative location of each of the conserved residues. Following from the above analysis, 10 residues were chosen for further study as they were located on the surface of the protein (Supplementary Fig S1B) and were either absolutely conserved throughout the alphaviruses, or in other cases were substituted by residues with similar physical characteristics (specifically the corresponding residue for both Met219 and Val260 in CHIKV is leucine in SINV) (Figs 1B and S1A). We chose to make two single substitutions (nsP3 amino acid numbering: M219A and E225A), and four double substitutions of adjacent or closely located residues (R243A/K245A, P247A/V248A, V260A/P261A and C262A/C264A). The latter two residues were shown to be involved in zinc coordination in the SINV AUD domain [14]. These mutations were cloned into a CHIKV subgenomic replicon (CHIKV-D-Luc-SGR) (Fig. 1C), derived from the ECSA strain (ICRES) (kind gift from Andres Merits, University of Tartu). This construct contains two luciferase reporter genes, a renilla luciferase (RLuc) is fused in frame within the C-terminal hypervariable domain of nsP3 in ORF1 and a firefly luciferase (FLuc) replaces the structural protein encoding region of ORF2, allowing simultaneous assessment of both input translation and genome replication. As a negative control we also created a polymerase-inactive mutant (GDD-GAA in the active site of nsP4).

### CHIKV subgenomic replicons exhibited different phenotypes in human, mammalian and mosquito cells

To analyse the effects of the AUD mutations on CHIKV genome replication, the panel of mutant CHIKV-D-Luc-SGR RNAs were transfected into a range of cell lines. As both liver and muscle are target organs for CHIKV infection we used three human cell lines. The human hepatoma cell line Huh7 are well characterised and we have previously shown [15] that they efficiently support CHIKV replication. To test any potential role of the AUD in protecting CHIKV from innate immune sensing we also used the Huh7 derivative cell line Huh7.5, which have a defect in innate immunity due to a mutation in one allele of the retinoic acid-inducible gene I (RIG-I) [16]. To investigate the role of the AUD in infection of muscle cells we used a human rhabdomyosarcoma cell line, RD. Additionally, two other mammalian (non-human) cell lines were used: C2C12 (a murine myoblast cell line) and BHK-21 (baby hamster kidney cells) based on their ability to support high levels of CHIKV replication [15]. Lastly, we used two mosquito (Ae. *albopictus)* derived cell lines: U4.4 and C6/36. Of note C6/36 have a defect in RNA interference (RNAi) due to a frameshift mutation in the Dcr2 gene, leading to production of a truncated and inactive Dicer-2 protein [17]. Again, use of these cells was intended to allow us to assess any role of the AUD in counteracting mosquito innate immunity.

We first tested replication in the human hepatoma cell line, Huh7 (Fig. 2A). Wildtype CHIKV-D-Luc-SGR exhibited robust replication in these cells with FLuc levels (a measure of genome replication) increasing approx. 30-fold between 4-12 h post-transfection. Consistent with this, RLuc levels (reflecting both input translation and replication) increased between 4-12 h but then declined at 24 h to input levels, possibly due to preferential transcription of the sub-genomic RNA at later times. Of the mutants M219A exhibited a modest, non-significant, reduction in replication, E225A replicated as wild type, but the other four nsP3 mutants and the nsP4 GAA mutant failed to replicate. The replication defect was indicated by either a reduction or a minimal increase in both RLuc and FLuc values from 4-24 h. A similar picture emerged when the mutant panel was screened in Huh7.5 cells (Fig. 2B). Consistent with the defect in cytosolic RNA sensing, replication of wildtype, M219A and E225A was higher in Huh7.5 cells compared to Huh7, however, this did not allow replication of the inactive mutants. For RD human rhabdomyosarcoma cells, a slightly different picture emerged (Fig. 2C): firstly both M219A and E225A replicated to a similar level as wildtype. Secondly, the P247A/V248A mutant, which was unable to replicate in Huh7 or Huh7.5 cells, was able to replicate to a low level in RD cells. The other 3 mutants and nsP4 GAA again failed to replicate.

**Figure 2.**
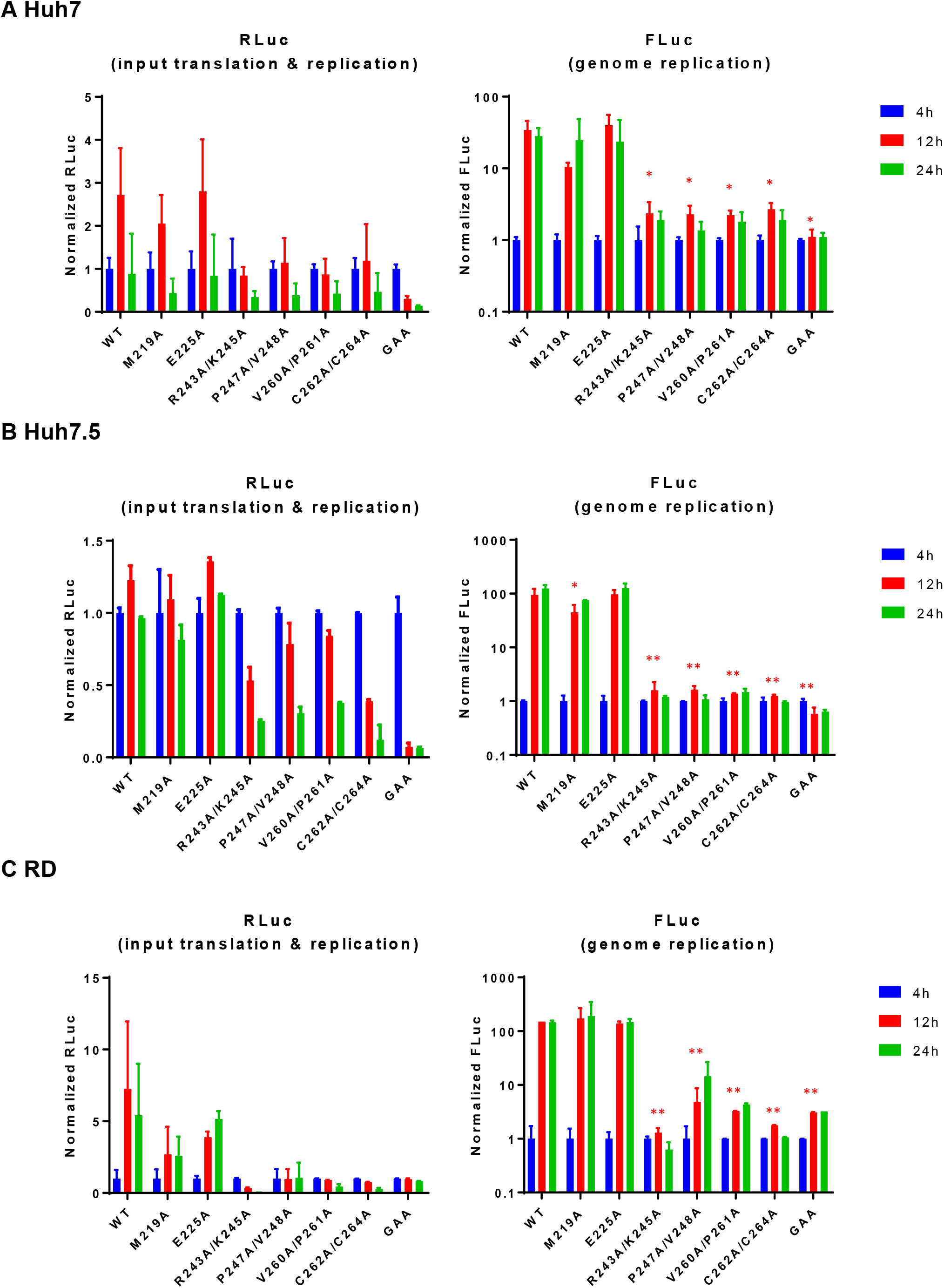
CHIKV AUD mutant replication in human cells. The indicated cells were transfected with CHIKV-D-luc-SGR wildtype and mutant RNAs and harvested for both RLuc and FLuc assays at the indicated time points. Luciferase values of wildtype and each mutant were normalized to 4 h values. (GAA: inactive mutant of nsP4 polymerase). Significant differences denoted by * (P<0.05), and ** (P<0.01), compared to wildtype.

We then evaluated the mutant panel in two other mammalian cell lines: C2C12 murine myoblasts (Fig 3A) and BHK-21 (Fig 3B). Wildtype CHIKV-D-Luc-SGR replicated to very high levels in both cell lines, with FLuc levels increasing ~1000-fold between 4-24 h. Overall the phenotypes of the panel of mutants were similar to those observed in RD, however two noticeable differences were observed. Firstly, R243A/K245A showed a low replication level in C2C12 cells. Secondly, P247A/V248A was capable of robust replication in both (albeit nearly 10-fold lower than wildtype). Interestingly, although FLuc levels for P247A/V248A were reduced, the concomitant RLuc levels were higher than wildtype, suggesting that although this mutant is replication competent there may be a defect in translation of ORF2. These data suggested that P247 and V248 were required for CHIKV genome replication in liver-derived cells, whilst enhancing but not essential for replication in cells derived from muscle or kidney, implying some cell type specific interactions of nsP3. V261A/P261A (adjacent to the zinc–binding site), and the zinc-coordinating cysteine mutant C262A/C264A were unable to replicate in either cell line, being indistinguishable from the GAA nsP4 control.

**Figure 3.**
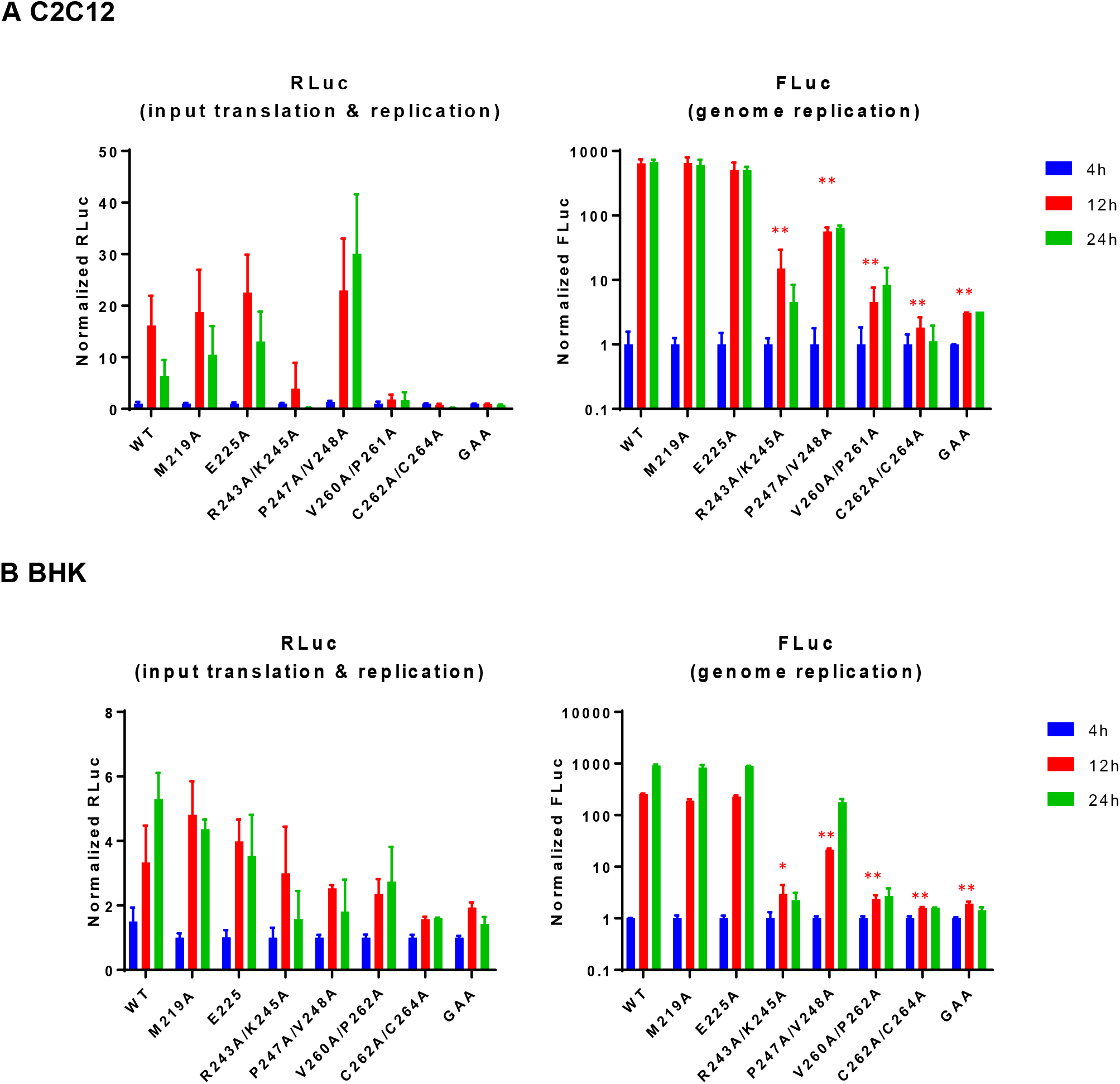
CHIKV AUD mutant replication in murine cells. The indicated cells were transfected with CHIKV-D-luc-SGR wildtype and mutant RNAs and harvested for both RLuc and FLuc assays at the indicated time points. Luciferase values of wildtype and each mutant were normalized to 4 h values. (GAA: inactive mutant of nsP4 polymerase). Significant differences denoted by * (P<0.05), and ** (P<0.01), compared to wildtype.

As a mosquito transmitted virus, CHIKV must replicate in both mammalian and mosquito cells. We therefore proceeded to evaluate the replicative capacity of the mutant panel in cells derived from the *Ae. albopictus* mosquito. Two cell lines were used: U4.4 and C6/36. The major difference between these two cell lines is that C6/36 have a defect in the RNAi response due to a frameshift mutation in the Dcr2 gene [17]. Consistent with this, although both mosquito cell lines supported robust replication, C6/36 supported higher levels than U4.4 (up to 1000-fold increase at 48 h). As described below, we observed remarkable differences in the mutant phenotypes in these cells compared to the mammalian cells (Fig. 4). The first difference was that M219A failed to replicate in U4.4 cells (Fig. 4A) but exhibited wildtype levels of replication in C6/36 cells (Fig. 4B), suggesting that M219 might be involved in interacting with, and inhibiting, the mosquito cell RNAi pathway. Secondly, R243A/K245A, which was unable to replicate in human cell lines and only showed low replication level in the C2C12 cells, was fully replication competent in both mosquito cell lines. Mutant P247A/V248A was partially replication competent in both cell lines, whereas as seen in mammalian cell lines neither V261A/P261A nor C262A/C264A replicated in mosquito cells.

**Figure 4.**
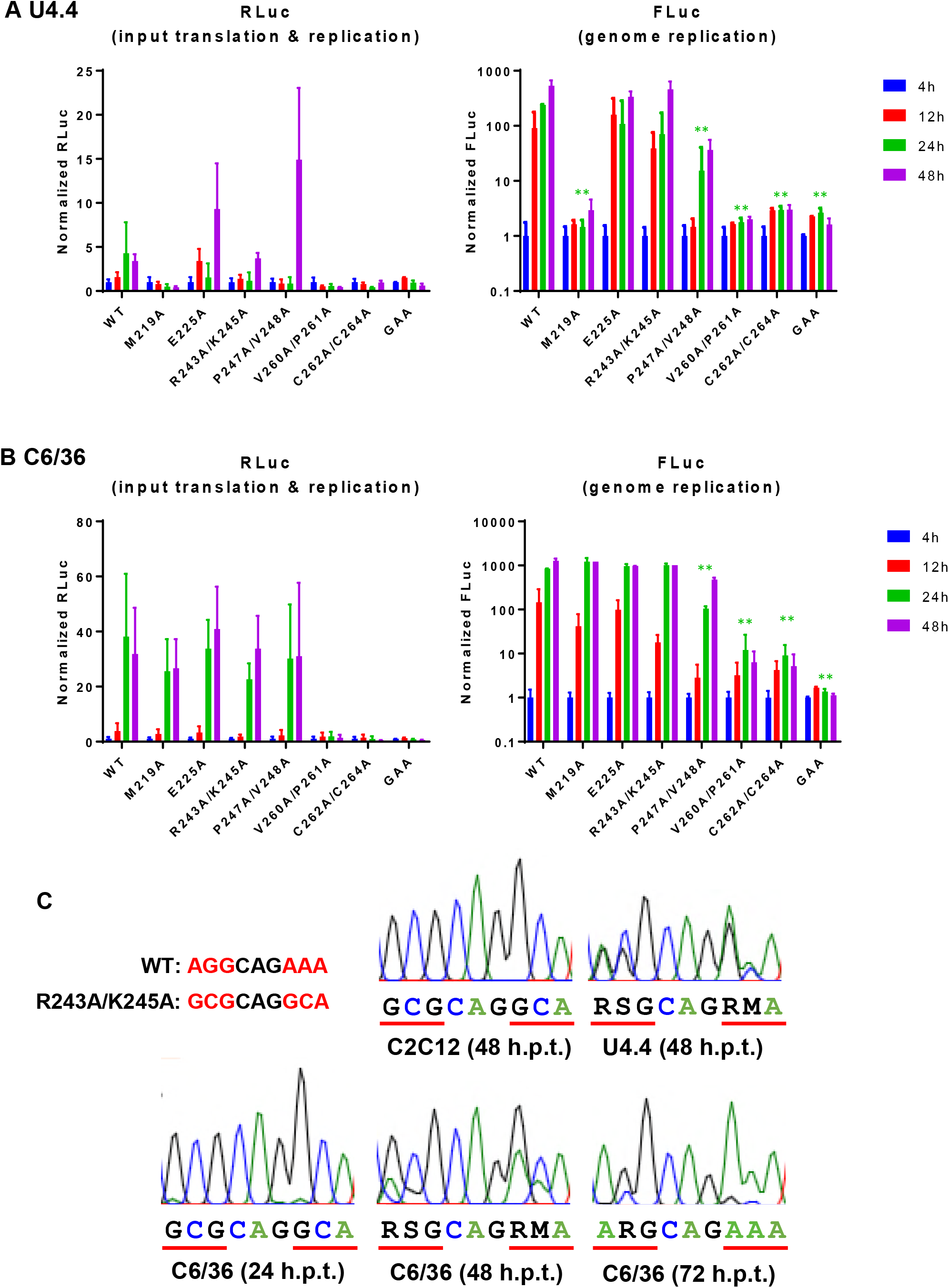
CHIKV AUD mutant replication in *Aedes albopictus* mosquito cells. (A, B) The indicated cells were transfected with CHIKV-D-luc-SGR wildtype and mutant RNAs and harvested for both renilla and firefly luciferase assay at the indicated time points. Luciferase values of wildtype and each mutant were normalized to 4 h values. (GAA: inactive mutant of nsP4 polymerase). Significant differences denoted by ** (P<0.01), compared to wildtype. (C) RT-PCR and sequencing analysis of CHIKV-D-luc-SGR-R243A/K245A. RNA was harvested at the indicated times, amplified by RT-PCR and sequenced. The wildtype and mutated sequences are shown below the sequence traces for reference. Nucleotide ambiguity codes used: R (A/G), S (G/C) and M (A/C).

The striking phenotypic difference between mammalian and mosquito cell lines for R243A/K245A led us to investigate this further. We considered that a simple explanation might be that the mutations had reverted in mosquito cell lines. To test this we extracted cytoplasmic RNA from C2C12, U4.4 and C6/36 cells at various time post transfection, and subjected them to RT-PCR and sequence analysis. In C2C12 cells at 48 h.p.t. we did not observe any sign of reversion (Fig 4C) – the sequence remained the same as the input RNA. However, for both U4.4 at 48 h.p.t., and C6/36 samples the sequence traces revealed the presence of a mixed population of mutant and wildtype. Notably we observed a sequential accumulation of revertants in the C6/36 samples: At 24 h.p.t. a very low proportion of revertants at the first position in the two codons was seen, at 48 h.p.t. the proportion increased and at 72 h.p.t. the sequences were almost entirely wildtype. These data are consistent with a requirement for R243 and K245 for efficient CHIKV genome replication.

### Role of the AUD in the context of infection of mammalian cells with CHIKV

We then sought to determine if the AUD played any role in other stages of the virus lifecycle. To test this, a subset of mutations that were able to replicate in all, or some of, the mammalian cells tested (M219A, E225A and P247A/V248A), together with the nsP4 GAA mutant as negative control, were introduced into an infectious CHIKV construct (ICRES-CHIKV). Capped *in vitro* transcribed virus RNA was electroporated into C2C12 cells and production of infectious virus was assessed by plaque assay of cell supernatants at 8, 24 and 48 h.p.e. C2C12 cells were chosen as our previous analysis had revealed that CHIKV grew to very high titres in these cells [15], and they are physiologically relevant, being muscle derived. As expected (Fig 5A), wildtype CHIKV produced a high titre of infectious virus following electroporation of C2C12 cells whereas the nsP4 GAA mutant did not produce any infectious virus. M219A and E225A were indistinguishable from wildtype, but P247A/V248A showed a significantly lower titre (approx. 10-fold reduced). Importantly, we showed by RT-PCR and sequence analysis of the infectious virus stocks, that they all retained the mutant sequence and had not undergone reversion to wildtype (Supplementary Fig S2).

**Figure 5.**
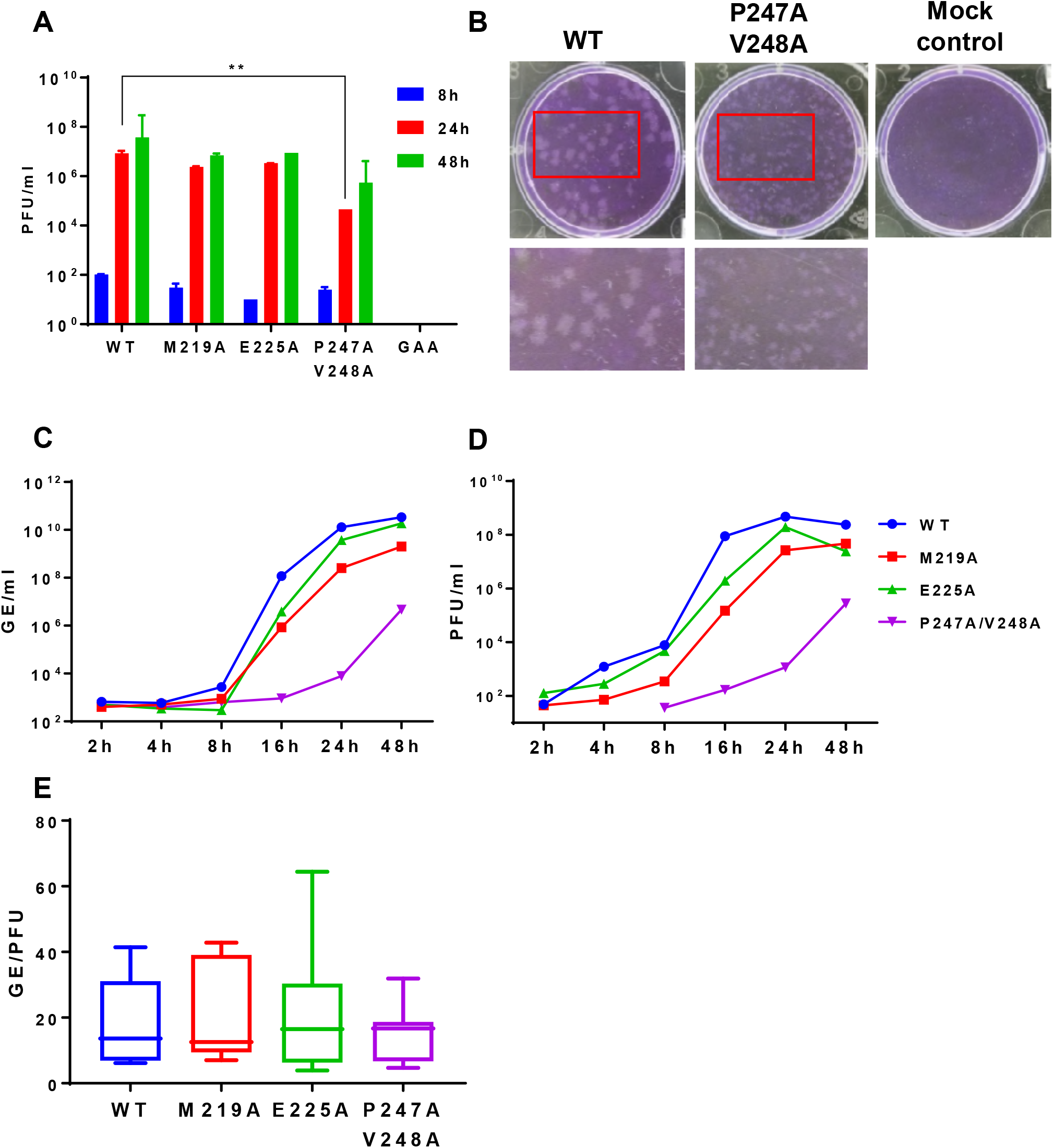
Phenotype of AUD mutations in the production of infectious virus. (A) ICRES-RNAs were electroporated into C2C12 cells and supernatants were collected at 48 h.p.e. Virus was titrated by plaque assay in BHK-21 cells. (B) Plaques for wildtype and P247A/V248A were visualised illustrating the small plaque phenotype for this mutant. (C-E) C2C12 cells were infected with CHIKV (wildtype and mutants) at an MOI of 0.1. Supernatants were collected at the indicated times for genome RNA quantification (qRT-PCR) (C) and virus titration by plaque assay (D). The ratios of genome RNA:infectivity were determined from (C) and (D) at 16, 24 and 48 h.p.i. and presented graphically (E).

During the course of these assays we noted that the P247A/V248A mutant uniquely exhibited a much smaller plaque size than the wildtype (Fig. 5B). We reasoned that this might reflect a defect in either virus production or spread that could be masked in the electroporation procedure due to the high level of input RNA. To test this we performed a one-step growth assay by infecting C2C12 cells at an MOI of 0.1 with either wildtype CHIKV or the 3 mutants (M219A, E225A and P247A/V248A). Cell supernatants were harvested at various times post infection and analysed for genomic RNA by qRT-PCR (Fig 5C, and infectivity by plaque assay (Fig 5D). Wildtype, M219A and E225A showed a rapid increase in both genomic RNA and infectivity between 8-48 h.p.i., reaching very high titres (for wildtype: 3.4×10^10^ RNA copies/ml and 4.7×10^8^ pfu/ml). In contrast, levels of P247A/V248A accumulated very slowly, reaching a maximum of 4.6×10^6^ RNA copies/ml and 2.8×10^5^ pfu/ml at 48 h.p.i. However, direct comparison of the genomic RNA quantification with the infectivity revealed that the specific infectivity of all four viruses were indistinguishable (Fig 5E). We conclude that, although P247A/V248A exhibited a defect in production of virus particles, the virions produced were equally infectious as wildtype.

We then asked whether the reduced virus titre exhibited by P247A/V248A resulted from a defect in virus assembly or release from infected cells. To address this question we first analysed levels of viral genomic RNA (by qRT-PCR) and infectious virus (by plaque assay) present at 24 h.p.i. within cells infected with wildtype or the 3 mutants (at an MOI of 1). This analysis (Fig 6A) revealed that the levels of intracellular genomic RNA for wildtype, M219A and E225A were comparable whereas P247A/V248A showed a 1000-fold reduction. This was consistent with the replicon data – for P247A/V248A genomic RNA levels were reduced from a mean of 7.9 × 10^9^ RNA copies/ml to 7.4×10^6^. Levels of infectious virus were similar for wildtype, M219A and E225A, however, uniquely P247A/V248A showed a dramatic 10^7^-fold reduction in the amount of intracellular infectious virus compared to wildtype: from 2.4 × 10^8^ pfu/ml to 9.8 × 10^1^ pfu/ml. This was reflected in a dramatic change in the ratio of genomic RNA:infectivity (Fig 6B), suggesting that although P247A/V248A exhibited a defect in genome replication there was an additional, more substantial phenotype in the production of infectious virus particles. The difference in magnitude of these two phenotypes suggests that they represent different functions of the AUD. To provide further support for this observation, we performed a similar experiment in which C2C12 cells were electroporated with wildtype or the 3 mutant virus RNAs (Fig 6C, D). This analysis revealed that the ratio of extra-to intracellular virus titres was significantly higher for P247A/V248A compared to wildtype and the other two mutants. We conclude that, although P247A/V248A produces less infectious virus, this can be released from the infected cells more efficiently than wildtype and the other two mutants.

**Figure 6.**
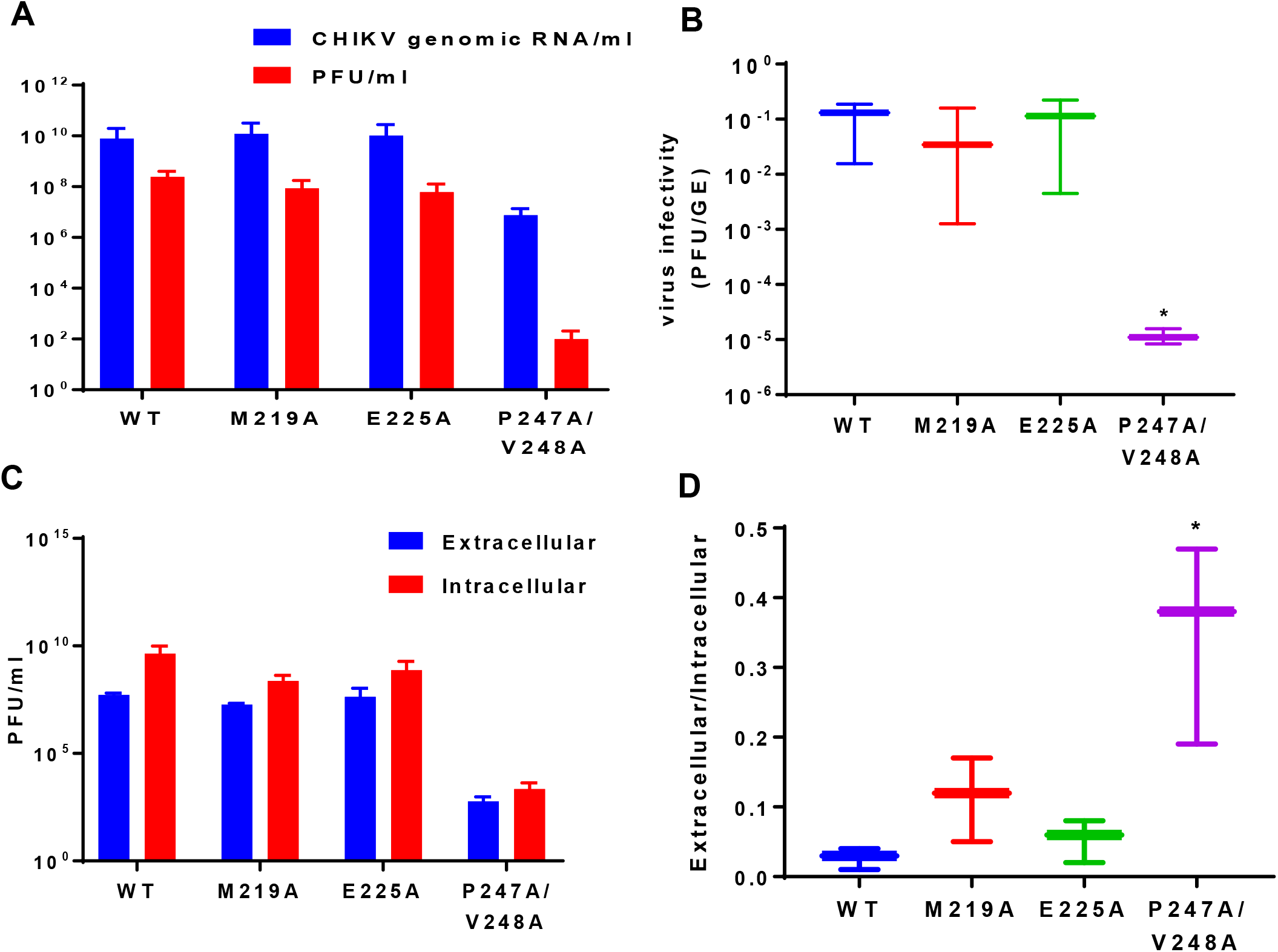
Phenotype of AUD mutations on virus entry, release and assembly. (A) C2C12 cells were infected with CHIKV at MOI of 1. At 24 h.p.i, cells were washed with PBS and resuspended in 1 ml fresh medium. Cell suspensions were freeze/thawed 3 times to release intracellular virus. Genome RNA was quantified by qRT-PCR, and virus titrated by plaque assay. (B) Graphical representation of the ratio of infectivity to genomic RNA. (C) Intracellular and extracellular viruses were collected at 36 h.p.e from C2C12 cells electroporated with the indicated ICRES RNA, and titrated by plaque assay. (D) Graphical representation of the ratio of extracellular to intracellular virus titres. Significant difference denoted by * (P<0.05) compared to wildtype.

### The P247A/V248A mutation selectively impairs subgenomic RNA synthesis

We considered that the reason for the reduction in virus assembly exhibited by P247A/V248A could be due to a direct role of nsP3 in this process, or some defect in the production of the structural proteins. To test this we analysed electroporated cells by western blot for the presence of both nsP3 and the capsid protein. P247A/V248A exhibited a modest reduction in nsP3 expression but a much greater reduction in the level of capsid expression. Indeed the ratio of capsid to nsP3 expression determined from the western blot analysis was approximately 10-fold lower for P247A/V248A (Fig 7A). During alphavirus replication the non-structural proteins (including nsP3) are translated from the full-length genomic RNA (gRNA), whereas the capsid and other structural proteins are translated from a subgenomic RNA (sgRNA). Transcription of both positive sense RNAs is mediated by a complex of the four nsPs using the full-length negative strand as template. This complex either initiates transcription from the 3’end of the negative strand or from the sub-genomic promoter. We hypothesised that the reduction in capsid expression for P247A/V248A could result from a defect in sgRNA transcription. To test this, C2C12 cells were electroporated and treated with actinomycin D (ActD) to block cellular RNA synthesis, prior to labelling with [^3^H]-uridine. Cellular RNA was extracted and analysed by MOPS-formaldehyde gel electrophoresis and autoradiography. As shown in Fig 7B, for WT, M219A and E225A, 2 radiolabelled species corresponding to gRNA and sgRNA were detected. However, for P247A/V248A, the sgRNA was present at very low levels, almost undetectable. The corresponding ratio of gRNA:sgRNA for P247A/V248A (25.3:1) was significantly higher than that of WT (1.5:1). As controls, mock electroporated cells treated with ActD contained no [^3^H]-labelled RNA species, whereas in the absence of ActD the expected smear of [^3^H]-labelled RNAs with predominant bands corresponding to 18S and 28S ribosomal RNAs were observed (Fig 7B). To confirm these results, the harvested RNAs were also analysed by sucrose gradient centrifugation. Consistent with the electrophoretic analysis, wildtype, M219A and E225A showed two peaks corresponding to sgRNA and gRNA, whereas P247A/V248A exhibited a dramatically reduced sgRNA peak (Fig 7C).

**Figure 7.**
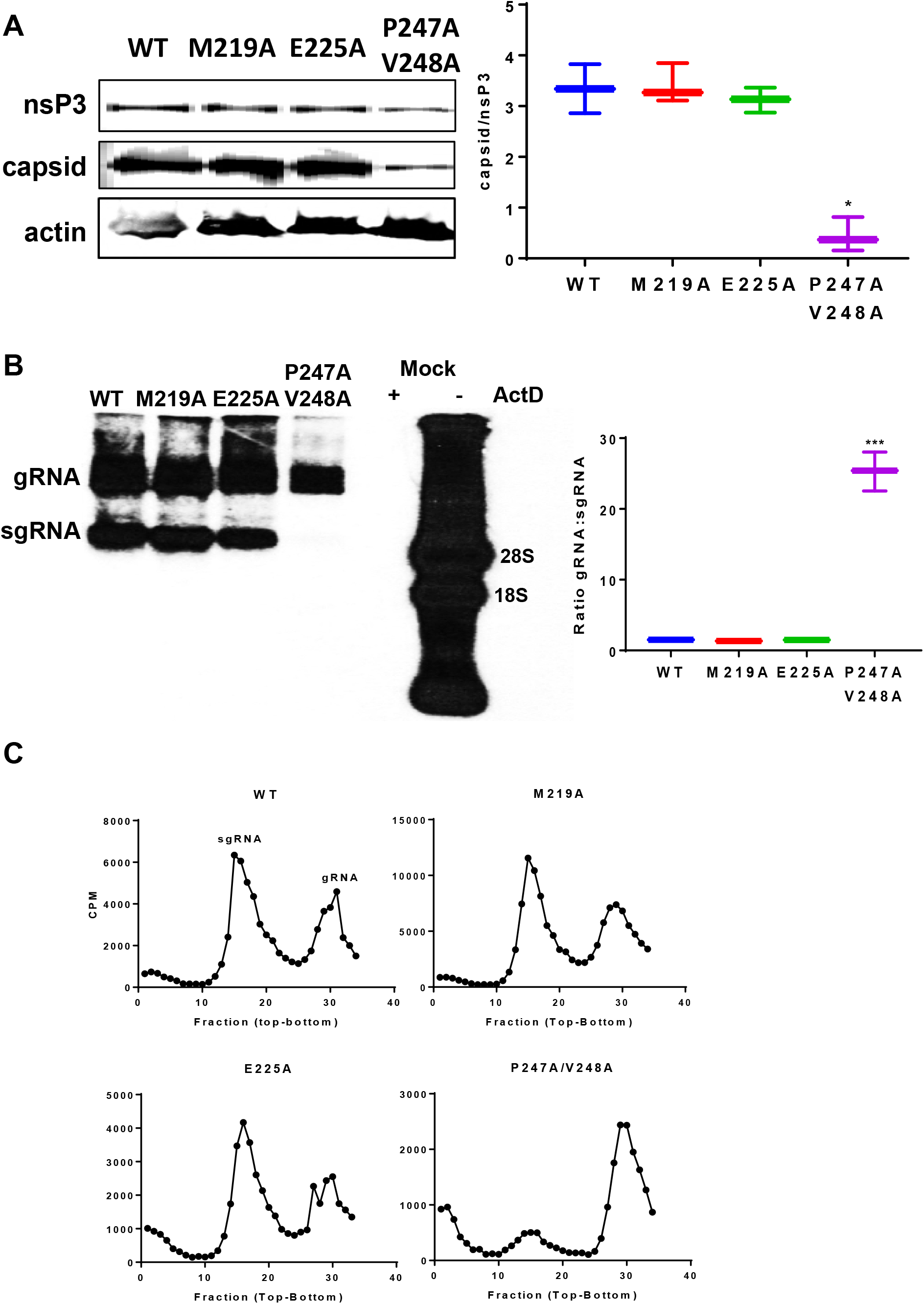
Effect of AUD mutations on CHIKV protein expression and RNA synthesis. (A) C2C12 cells were electroporated with ICRES-RNAs and cell lysates were collected at 36 h.p.e. Expression of nsP3 and capsid was analysed by western blot. Multiple western blots were quantified using a LiCor Odyssey Sa fluorescence imager and the graph on the right shows the ratio of capsid to nsP3 expression. (B) C2C12 cells were electroporated with the indicated ICRES RNA, cellular RNA synthesis was inhibited by actinomycin D and nascent viral RNAs were labelled with [^3^H]-uridine. The graph on the right shows the ratio of gRNA to sgRNA. (C) The same RNAs were fractionated on a sucrose gradient and [^3^H]-labelled RNAs were detected by scintillation counting of individual fractions.

### Effect of the P247A/V248A mutation on the RNA-binding activity of the AUD

Our data thus far are consistent with the hypothesis that the P247A/V248A mutation results in a reduction in the ability of the nsP complex to recognise and initiate transcription from the subgenomic promoter. As the AUD has been predicted to possess RNA-binding activity [14], and initiation of gRNA or sgRNA transcription by the nsP complex will require specific recognition of cognate RNA sequences on the CHIKV negative strand, we further postulated that the phenotype of the P247A/V248A might be explained by a defect in RNA binding activity. To test this we expressed wildtype and P247A/V248A AUD mutants in *E.coli* as His-Sumo fusion proteins. The AUDs were cleaved from the fusion proteins by Sumo-protease and analysed by SDS-PAGE. As shown in Fig 8A, the AUDs could be purified to a high degree of homogeneity. Circular dichroism (CD) analysis (Fig 8B) revealed that, as expected, the AUDs comprised predominantly α-helix with no significant differences in the overall structure as a result of the mutations. To test for RNA-binding activity we performed a filter binding assay [18] using purified AUDs and a radiolabelled RNA corresponding to the 3’ end of the CHIKV genome (3’UTR(+)), the 3’ end of the genomic negative strand RNA (5’UTR(-)), or the negative strand subgenomic promoter (sg-prom(-)) (Fig 8C). Both wildtype and P247A/V248A AUD were able to bind the 3’UTR(+) (Fig 8D) and 5’UTR(-) (Fig 8E), however P247A/V248A exhibited a significant increase in Kd values and decrease in maximal binding levels (endpoints) compared to wildtype. As 3’UTR(+) and 5’RNA(-) are involved in the initiation of negative and positive strand genome RNA synthesis, respectively; impaired binding of the P247A/V248A mutant AUD may explain the observed defect in CHIKV genome replication (Fig 3). For binding to the sg-prom(-) RNA (Fig 8F), P247A/V248A AUD showed a different phenotype with both a higher endpoint and Kd than wildtype. Kd and endpoint values are listed in Fig 8G. This result suggests that the P247A/V248A defect in sgRNA synthesis may be in part explained by a reduction in the ability to specifically bind the subgenomic promoter.

**Figure 8.**
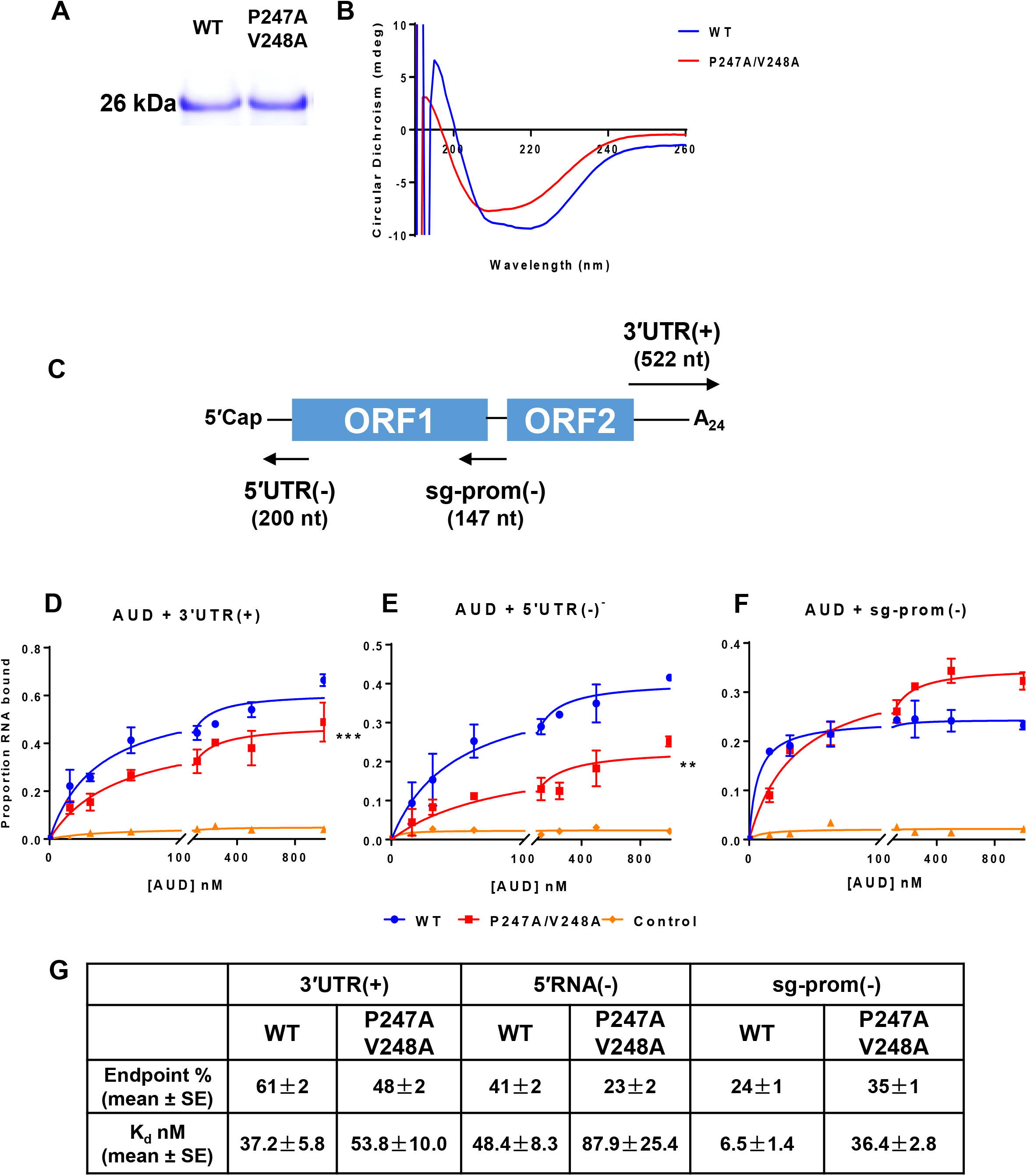
AUD RNA-binding activity to different viral RNAs. (A) *E. coli* expressed AUD (wildtype and P247A/V248A) analysed by SDS-PAGE and Coomassie blue staining. (B) Circular Dichroism analysis of purified AUD. (C) Schematic of the CHIKV genome showing the location of the various RNAs used in subsequent filter binding analysis. (D-F) Filter binding analysis of the interaction between AUD and the indicated RNA species. Purified AUD at the indicated concentrations was incubated with radiolabelled RNA (1 nM) before application to a slot blot apparatus, filtering through nitrocellulose (protein-RNA complex) and Hybond-N (free RNA) membranes, and visualization by phosphoimaging. The negative control is wildtype AUD with an 80-mer aptamer raised against the foot-and-mouth disease virus 3D RNA-dependent RNA polymerase [18]. The percentage of RNA bound to the nitrocellulose membrane was quantified and plotted as a function of the AUD concentration. The data was fitted to a hyperbolic equation. (G) Endpoint (% of total RNA bound) and Kd values derived from the graphs in (D-F).

To explore the RNA binding activity of nsP3 to CHIKV genomic RNA during virus replication, we exploited a previously generated derivative of the ICRES infectious clone in which a twin-strep tag (TST) was introduced in frame near the C-terminus of nsP3, allowing efficient affinity purification of nsP3 by streptactin chromatography. We had previously used this experimental approach to investigate protein-protein and protein-RNA interactions of the hepatitis C virus NS5A protein [19–21]. C2C12 cells were electroporated with either wildtype or P247A/V248A mutant TST-nsP3 CHIKV RNAs. NsP3 proteins were purified from cell lysates on streptactin beads and analysed by western blot for nsP3 (Fig 9A) and qRT-PCR to determine the amount of gRNA associated with nsP3 (Fig 9B). Consistent with the *in vitro* RNA filter binding assay data, P247A/V248A bound approximately 10-fold less gRNA compared to wildtype (Fig 9C).

**Figure 9.**
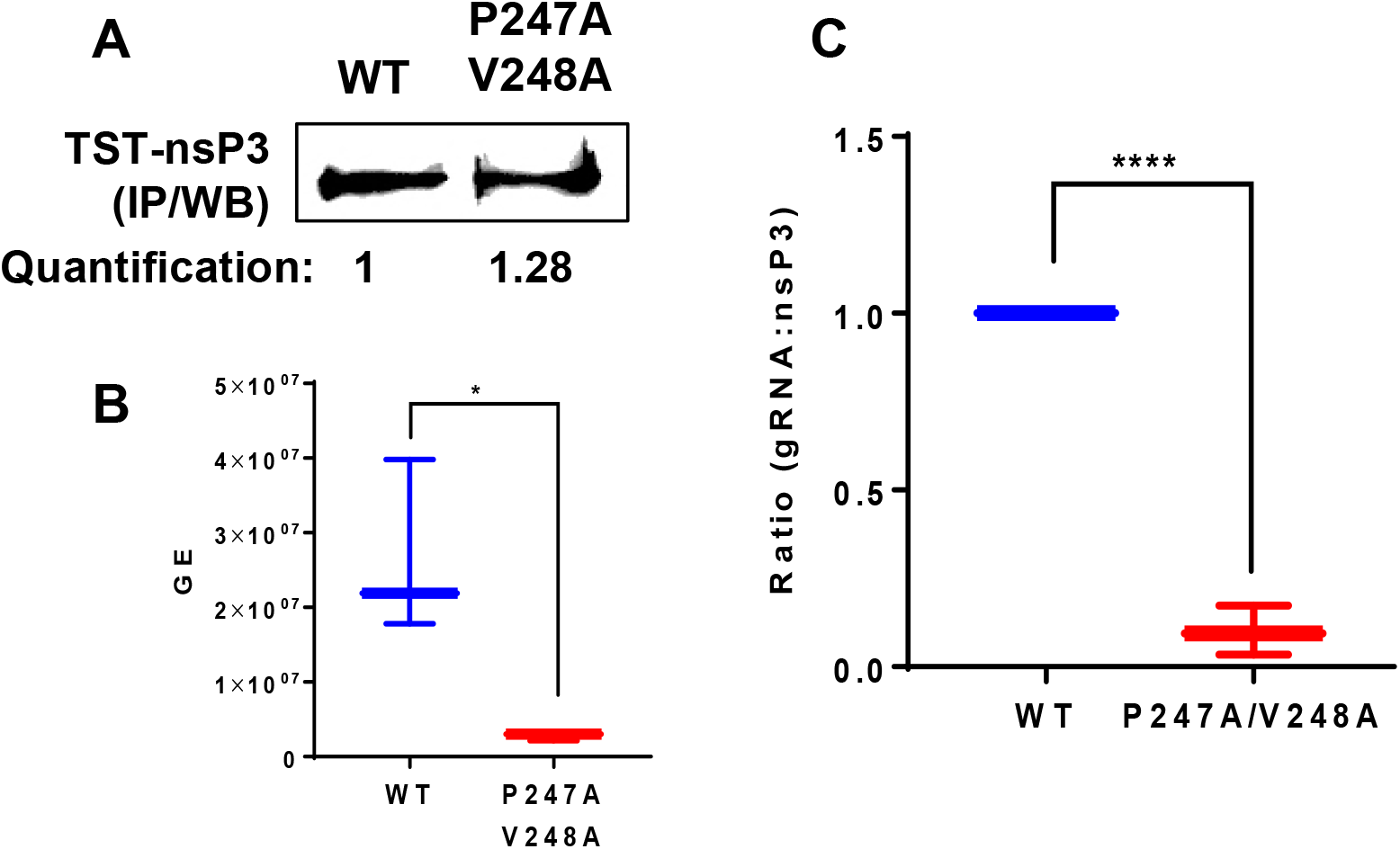
CHIKV genome RNA association with nsP3 during virus replication. C2C12 cells were electroporated with ICRES nsP3-TST RNAs. Cell lysates were collected at 60 h.p.e. and nsP3-TST was precipitated with Streptactin-sepharose beads. Bound proteins were subjected to western blotting (A) and co-precipitated RNAs were extracted by TRIzol and quantified by qRT-PCR (B). The ratio of gRNA to nsP3 is depicted graphically (C).

### Sub-cellular localisation of nsP3, capsid and dsRNA during CHIKV replication

The effect of P247A/V248A on the nsP3:gRNA interaction suggested that this mutation might also disrupt the subcellular localisation of nsP3 in relation to both replication complexes and sites of virion assembly. To test this we exploited another derivative of the ICRES infectious CHIKV clone in which ZsGreen was inserted into nsP3 at the same position as the TST tag [22, 23]. C2C12 cells were electroporated with ICRES-nsP3-ZsGreen-CHIKV RNAs (wildtype or P247A/V248A), and cells were analysed by confocal laser scanning microscopy (CLSM) with Airyscan for the distribution of nsP3, capsid (as a marker for virion assembly sites) and dsRNA (as a marker of genome replication) at different times post-electroporation. For wildtype at 4 h.p.e. (Fig 10), small clusters of nsP3, capsid and dsRNA appeared in the cytoplasm but there was little co-localisation. By 8 h.p.e., nsP3, capsid and dsRNA co-localised in larger clusters, these appeared to accumulate at the plasma membrane at 12 and 16 h.p.e., by which time the majority of nsP3, capsid and dsRNA were co-localised on plasma membrane. By 24 h.p.e., it was clear that the infection cycle was complete as there was a reduction in levels of nsP3, capsid and dsRNA. Interestingly, capsid and dsRNA were still co-localised at the plasma membrane while most nsP3 was perinuclear. In contrast, P247A/V248A exhibited a very different distribution pattern of all three markers throughout the infection cycle (Fig 11). Consistent with the western blot data (Fig 7A) levels of capsid and dsRNA were lower than wildtype at all timepoints, but in addition the co-localisation of nsP3, capsid and dsRNA was markedly reduced and the three markers never accumulated at the plasma membrane as seen for wildtype. Consistent with the delay in virus release shown in Fig 5, it was clear that, unlike wildtype, the infection cycle was not complete by 24 h.p.e. as levels of nsP3 and capsid were highest at this timepoint.

**Figure 10.**
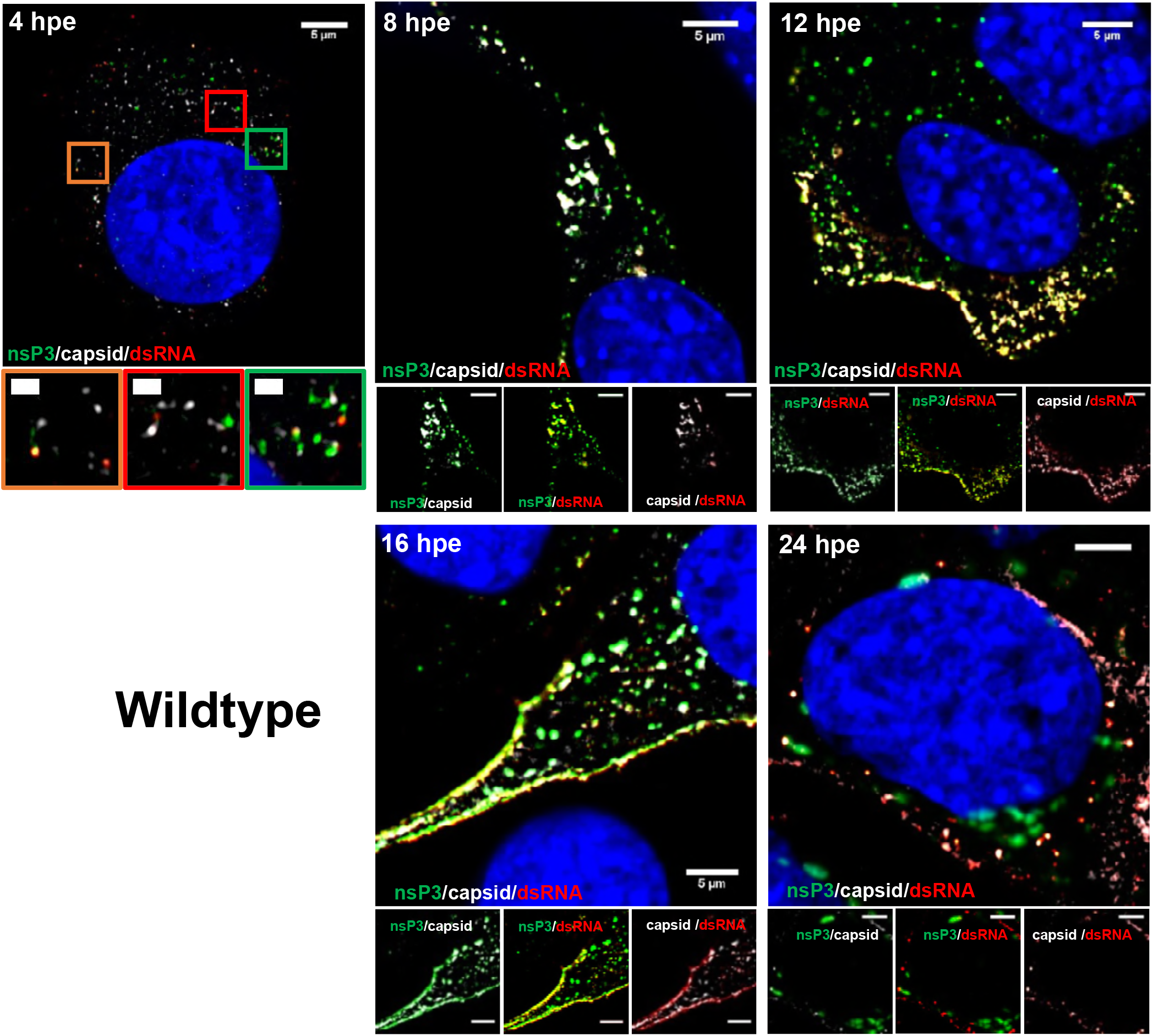
Fluorescence analysis of nsP3, capsid and dsRNA distribution during infection of C2C12 cells with wildtype CHIKV. C2C12 cells were electroporated with ICRES-nsP3-ZsGreen-CHIKV RNA. Cells were fixed at the indicated time points post-infection and stained with antibodies to capsid protein (white) and dsRNA (red). Green: nsP3-ZsGreen fusion, blue: nuclear DAPI counterstain. The scale bars are 5 μm and 1 μm, respectively.

To provide a quantitative assessment of the differences between wildtype and P247A/V248A we quantified the percentage co-localisation of nsP3 with either dsRNA (Fig 12A) or capsid (Fig 12B) from 5 cells at each timepoint. As shown in Fig 12A, at all timepoints the percentage co-localisation of nsP3 with dsRNA was significantly lower for P247A/V248A. The results for nsP3 co-localisation with capsid were less clear-cut: for wildtype there was a gradual increase from 8-24 h.p.e., however for P247A/V248A levels remained fairly constant with a transient drop at 16 h.p.e., consistent with the confocal images in Fig 11. These data are consistent with a role for the AUD, and residues P247/V248 in particular, in mediating interactions between nsP3 and both genome replication complexes and virus assembly sites. They also suggest that nsP3 is not only involved in virus genome replication, but may also play a role in the trafficking of capsid to plasma membrane.

**Figure 11.**
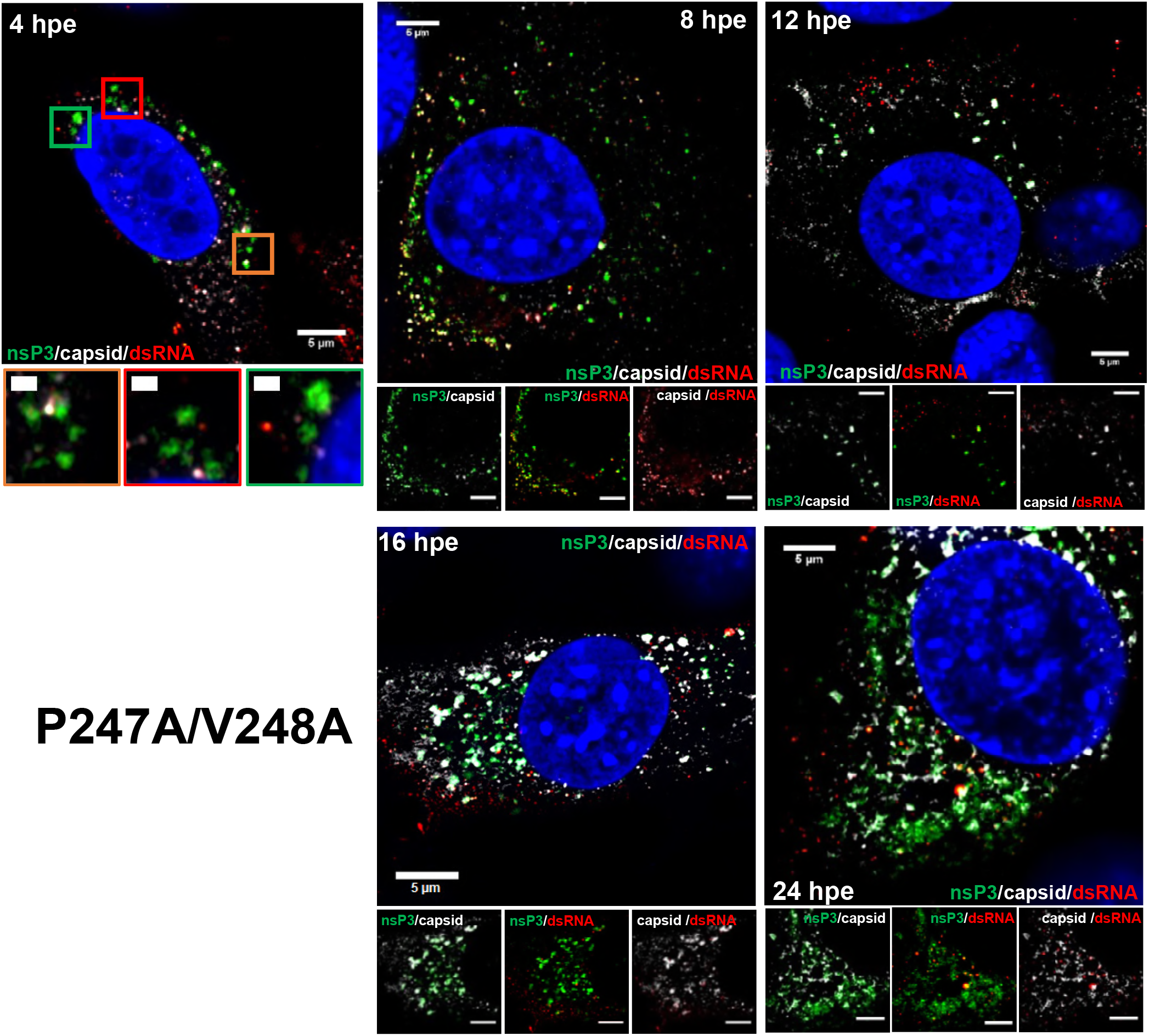
Fluorescence analysis of nsP3, capsid and dsRNA distribution during infection of C2C12 cells with P247A/V248A mutant CHIKV. C2C12 cells were electroporated with ICRES-nsP3-ZsGreen-CHIKV-P247A/V248A RNA. Cells were fixed at the indicated time points post-infection and stained with antibodies to capsid protein (white) and dsRNA (red). Green: nsP3-ZsGreen fusion, blue: nuclear DAPI counterstain. The scale bars are 5 μm and 1 μm, respectively.

**Figure 12.**
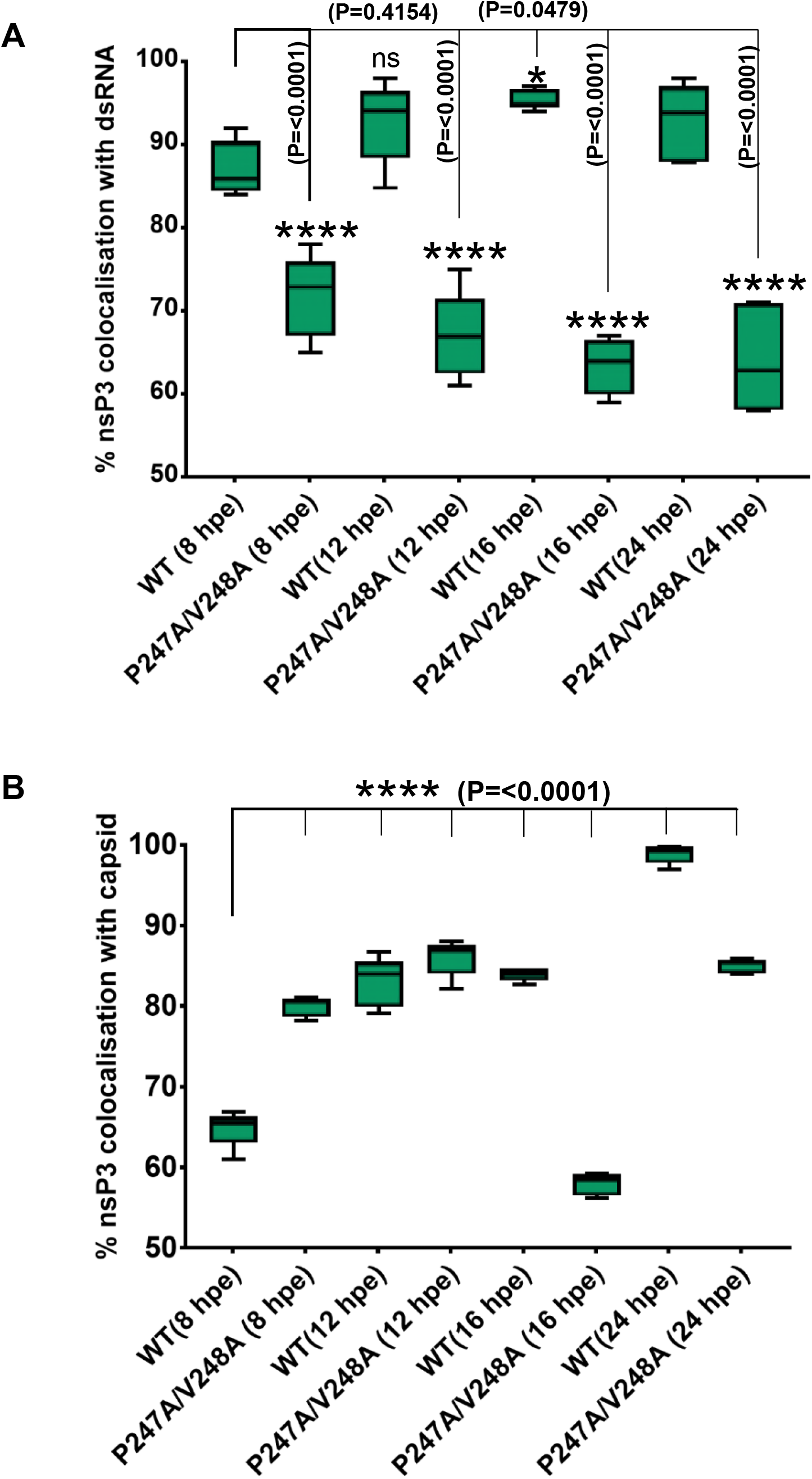
Co-localisation of nsP3 with capsid protein and dsRNA during virus replication. (A) Quantification of the percentages of nsP3 colocalised with dsRNA. Co-localisation of nsP3 with dsRNA. (B) Quantification of the percentages of nsP3 colocalised with capsid. Co-localisation analysis (green blocks) were determined from 5 cells for each construct using Fiji. ****, * indicates significant difference (P<0.0001, P<0.0479, respectively) from the results for WT.

## Discussion

Of the four alphavirus non-structural proteins, nsP3 remains the least well understood [24]. The protein consists of three domains, the N-terminal of which has been identified as a macrodomain that binds to ADP-ribose and possesses ADP-ribosylhydrolase activity [9]. Recent studies have proposed a role for this enzymatic activity in virus pathogenesis but as yet the underlying mechanisms remain elusive [11]. The C-terminal hypervariable domain differs dramatically in amino acid sequence between different alphaviruses and is intrinsically disordered. It has been shown to interact with a range of cellular proteins, including components of stress granules [25], and is implicated in the assembly of virus genome replication complexes.

In contrast, we know virtually nothing about the function of the central AUD domain. The fact that this domain is highly conserved between different alphaviruses suggests that it plays a fundamental role in the virus lifecycle. Detailed structural information about the AUD is available however, as the partial structure of the SINV nsP2-nsP3 precursor, including the C-terminal protease and methyltransferase-like domains of nsP2 and the macro and AUD domains of nsP3, has been determined [14]. This analysis revealed that the AUD contained an unique zinc-binding fold with four cysteine residues coordinating a zinc molecule, this formed part of a putative RNA binding surface. Mutagenesis of two of these cysteines revealed an essential role in virus replication. Our data agree with this observation, as the C262A/C264A mutant failed to replicate in any cell type tested.

Mutation of two residues adjacent to the zinc-binding cysteines, V260A/P261A, also completely abrogated CHIKV genome replication. Although adjacent in the primary amino acid sequence, these residues are located on the distal face of the AUD (Fig 1B), suggesting that they are not involved in zinc binding, but may instead interact with key cellular factor(s) or play an alternative structural role.

In contrast, the other mutants generated during this study exhibited a number of distinct cell-type and species-specific phenotypes (summarised in Table 1). Mutation of two surface exposed basic residues (R243 and K245) abrogated replication in all mammalian cells but showed full replication capability in mosquito cells. However, this apparent discrepancy could be explained by the observation that these two mutations rapidly reverted to wildtype in mosquito cells but failed to do so in C2C12 cells (Fig 4C). These data indicate that R243 and K245 are required for CHIKV genome replication. We do not have an explanation for why R243A/K245A, but not the other lethal mutations, was able to revert in mosquito cells. However it is noteworthy that the reversion to the wildtype sequence took 72 h in C6/36 cells, suggesting that perhaps the lack of cytopathology of CHIKV replication in mosquito cells could facilitate the replication of a minority species. Interestingly, the sequence trace at 24 h in C6/36 shows the presence of a such minority species that would encode a Thr at 243 and 245, suggesting that the two basic residues are not absolutely required.

**Table 1:**
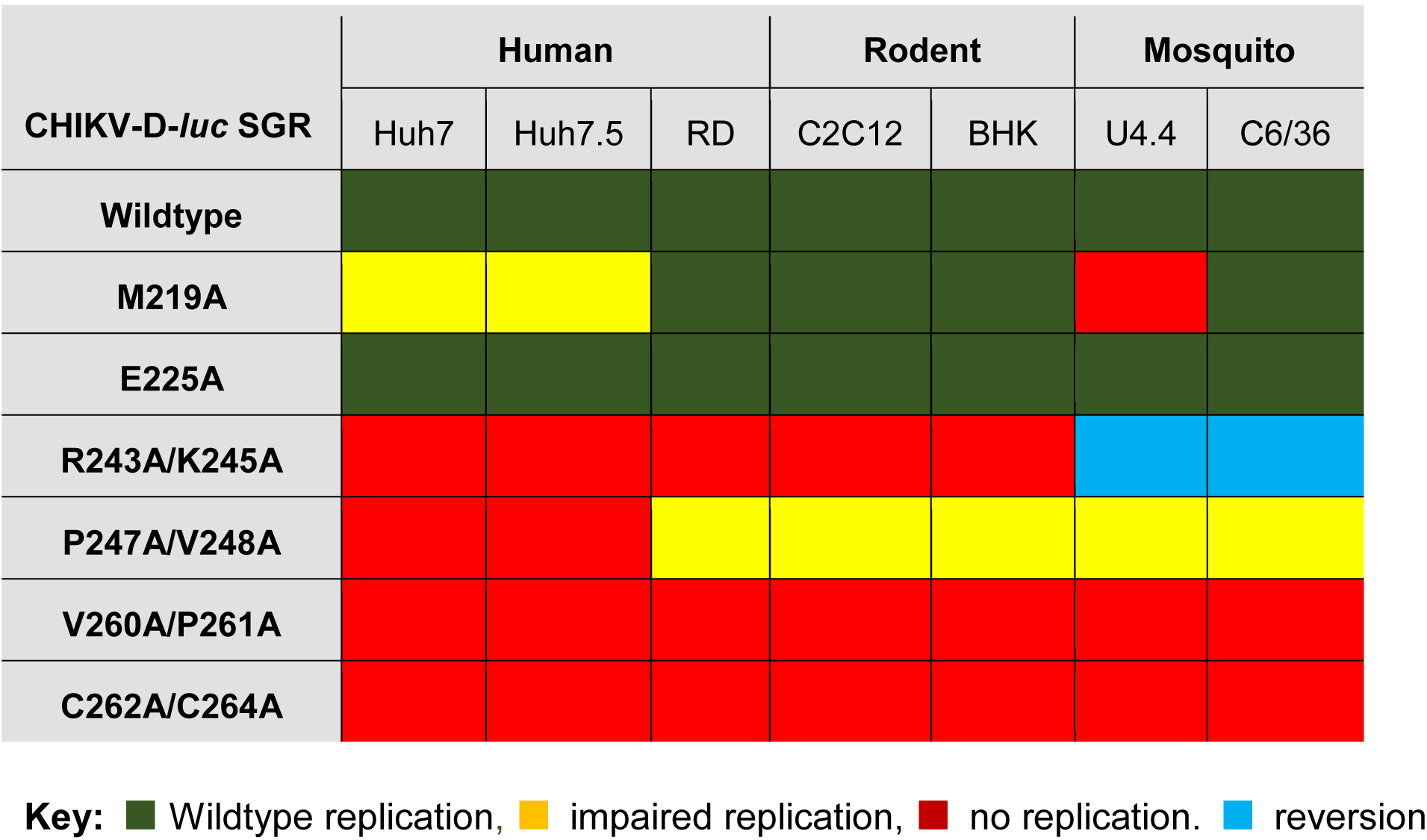
AUD mutant replication phenotypes in different cell types.

M219 was also of particular interest as mutation of this residue had no significant effect on genome replication in any cell type apart from U4.4 mosquito cells. M219A replicated well in C6/36 mosquito cells and the key difference between these two cell lines is that C6/36 have a defect in the RNAi response due to a Dcr2 mutation [17]. We propose therefore that M219 may interact with a component of the mosquito RNAi response to inhibit this key mosquito antiviral pathway. We are currently undertaking proteomic and functional analysis to test this hypothesis.

In the second part of this study we focussed on P247A/V248A to address the molecular mechanism underpinning the phenotype of this mutant. In the context of the subgenomic replicon P247A/V248A showed a variety of phenotypes from complete lack of replication in Huh7 and Huh7.5 cells (Fig 2), to a 10-fold reduction in other mammalian and mosquito cells (Fig 3). Virus data in C2C12 cells were consistent with replicon results: following electroporation of viral RNA P247A/V248A showed a modest but significant defect in virus infectivity and exhibited a small plaque phenotype. Further study indicated that P247A/V248A was competent in virus entry and virus release, however, it exhibited a major defect in assembly of infectious virus particles. This defect led to both a delay and a reduction in the release of infectious virus, consistent with the small plaque phenotype. The molecular mechanism underpinning the P247A/V248A defect was shown to be a reduction in subgenomic RNA synthesis, leading to a concomitant reduction in the expression of the structural proteins. It is noteworthy that when analysed in the context of the replicon in C2C12 cells, the P247A/V248A mutant exhibited a 10-fold reduction in FLuc, but RLuc was higher than wildtype. This is also consistent with a defect in transcription of the subgenomic mRNA. The reduced affinity of P247A/V248A AUD for the CHIKV sg-prom(-) RNA as demonstrated by the RNA filter binding assay may help to explain this, but it is possible that other effects of P247A/V248A, such as aberrant host protein recruitment, may also help to explain the defect in subgenomic RNA synthesis. Of note, P247A/V248A AUD also exhibited impaired binding to the 3’UTR(+) and 5’RNA(-) *in vitro*, and also bound less genomic RNA *in vivo*, compared to wildtype, consistent with an overall defect in RNA replication. Previous studies have also shown that nsP3 is important for initial replication complex formation and negative strand RNA synthesis [26]. Taken together we propose that the AUD plays a critical role in all stages of CHIKV RNA synthesis, but particularly in the transcription of the sgRNA.

Analysis of the distribution of nsP3, capsid and dsRNA during CHIKV replication by confocal microscopy revealed further insights into the P247A/V248A phenotype. Wildtype nsP3 exhibited a high level of co-localisation with dsRNA at all time points up to 24 h.p.e., consistent with the role of nsP3 in genome replication. At 12/16 h.p.e. both nsP3 and dsRNA also co-localised with capsid and were concentrated at the plasma membrane. In contrast, P247A/V248A nsP3 showed not only a significant reduction in the co-localisation with dsRNA, but also a loss of the plasma membrane accumulation. These observations are consistent with a role for nsP3 (and the AUD in particular) in coordinating the processes of genome replication and virus assembly to facilitate production of infectious virus particles at the plasma membrane. This is in agreement with early evidence for a juxtaposition of sites of genome replication, viral protein translation and nucleocapsid assembly in the case of Sindbis virus [27]. In addition it may be that nsP3 has a role in the trafficking of nucleocapsids from these sites (cytoplasmic vacuoles (CPVs)) to the plasma membrane.

In conclusion we propose that the nsP3 AUD is a multi-functional domain. It is not only a critical determinant of both cell and species-specificity, but also plays roles in virus genome replication and assembly. The rational mutagenesis of the CHIKV AUD described here is the first detailed structure-function analysis of this domain and raises many questions. In particular, we need to determine what cellular and viral proteins are interaction partners for the AUD, and investigate how these interactions shed light on the phenotypes of AUD mutants? We expect that our current proteomic analysis, exploiting both a novel SNAP-tagged nsP3 recently developed in our laboratory [28] and a One-Strep tag (OST) approach which we recently used to identify interacting partners of the hepatitis C virus NS5A protein [19], will provide some of the answers to these questions. We also hope that these studies will help to identify targets for antiviral intervention and means of rational attenuation for vaccine development.

## Acknowledgements

We thank Andres Merits (Institute of Technology, University of Tartu, Estonia) for the CHIKV subgenomic replicons, ICRES constructs and antibodies to nsP3 and capsid. We thank Raymond Li (University of Leeds) for the OST-tagged ICRES constructs, and Joseph Ward (University of Leeds) for help and advice with the ^3^H-labelling.

## Materials and Methods

### Sequence alignment

AUD amino acid sequences of different alphaviruses (Sindbis, Ockelbo, O’Nyong-Nyong, Ross River, Semliki Forest, Fort Morgan, Venezuelan, Eastern and Western Equine Encephalitis, Highlands-J and CHIKV) were obtained from NCBI and aligned by Clustal Omega. The predicted locations of conserved residues were then identified by Pymol, taking the Sindbis nsP2/3 protein structure (PDB ID code 4GUA) [14] as a reference.

### Cell culture

Mammalian cells Huh7, Huh7.5, RD, C2C12 and BHK-21 were maintained at 37°C with 5% CO_2_ in DMEM supplemented with 10% FCS, 0.5 mM non-essential amino acids and penicillin-streptomycin (100 units/mL). Huh7 cells were obtained from John McLauchlan (Centre for Virus Research, Glasgow), Huh7.5 cells from Charles Rice (Rockefeller University, New York), RD cells from Nicola Stonehouse (University of Leeds), C2C12 cells from Michelle Peckham (University of Leeds) and BHK-21 cells from John Barr (University of Leeds). Mosquito cell lines (U4.4 and C6/36) were obtained from Susan Jacobs (The Pirbright Institute) and maintained in Leibovitz’s L-15 media supplemented with 10% FBS, 10% tryptose phosphate broth and penicillin-streptomycin (100 units/mL). Mosquito cells were incubated at 28°C without CO_2_.

### Construction of CHIKV subgenomic replicons and infectious viruses with AUD mutations

A fragment including the AUD was excised from the CHIKV-D-Luc-SGR plasmid, and inserted into pcDNA3.1 to generate pcDNA3.1-AUD. This was used as a template for site-directed (Quikchange) mutagenesis using specific primers (primer sequences available upon request) to produce the required AUD mutations. Finally, the AUD mutated fragments were excised and religated into the CHIKV-D-Luc-SGR. AUD fragments were subsequently excised from CHIKV-D-Luc-SGRs and ligated into ICRES-CHIKV-WT. For the One-Strep tag (OST) derivatives of wildtype and P247A/V248A synthetic oligonucleotides (sequence available upon request) were used to substitute the appropriate coding sequence [19] for the RLuc in nsP3.

### Transfection and Dual-luciferase Assay

Capped RNAs were generated from the CHIKV-D-Luc-SGR for transfection using the mMACHINE SP6 transcription kit (ThermoFisher Scientific) and purified with PureLink RNA Mini Kit (Life Technologies). Transfection of the CHIKV-D-Luc-SGR RNAs was performed in different cells using Lipofectamine 2000 (Life Technologies) according to the manufacturer’s instructions. At 4, 12, 24 and 48 h post transfection (h.p.t.), cells were harvested and both Renilla and Firefly luciferase activity measured using the Dual-luciferase Assay System (Promega) according to the manufacturer’s instructions. Each sample had three repeats and the data shown in this study represent the mean of three experimental replicates.

### Sequence analysis of subgenomic or viral RNA

Cytoplasmic RNA from electroporated or infected cells were Trizol extracted, prior to reverse transcription with random primers using SuperScript IV Reverse Transcriptase (Invitrogen) according to the manufacturer’s instructions. cDNAs were then used as a template to amplify part of nsP3 sequence (macro domain and AUD) with specific primers (primer sequences available upon request). PCR products were subjected to sequencing analysis.

### Titration of infectious CHIKV by plaque assay

ICRES-CHIKV RNAs were produced and purified as described above. C2C12 cells were electroporated with ICRES-CHIKV RNAs and incubated at 37°C. Cell supernatants were collected at 8, 24 and 48 hours post electroporation (h.p.e.), diluted with cell medium and applied to monolayers of BHK-21 cells for 1 h at 37°C. The inoculum was aspirated and plates were overlaid with 0.8% methylcellulose for 48 h at 37°C for plaque formation. For titration of intracellular viruses, cells were freeze/thawed 3 times and supernatants were collected by centrifugation at 12000×g for 10 min., prior to application to BHK-21 cells All virus work was performed in a Biological Containment Level 3 (BSL3) laboratory. Plaques were visualised by photography with a Canon EOS 80D.

### One-step virus growth curve

Infectious CHIKV was harvested from C2C12 cells electroporated with ICRES RNAs at 48 h.p.e. Cell supernatants were titrated by plaque assay in BHK-21 cells and stored at −80°C. For virus growth kinetic analysis, C2C12 cells were infected with wildtype or AUD mutant CHIKV at an MOI of 0.1 for 1 h. Infected cells were washed three times with PBS and incubated with fresh complete medium at 37°C. For RNA quantification, RNAs were extracted from supernatants Trizol (ThermoFisher Scientific). qRT-PCR for CHIKV genome RNA was performed with One-step MESA GREEN qRT-PCR MasterMix Plus for SYBR assay (Eurogentec) following the manufacturer’s instructions. Primer sequences available upon request. Plaque assay was performed as described above.

### CHIKV RNA synthesis

C2C12 cells were electroporated with ICRES RNAs for 10 h. Actinomycin D (1 μg/ml) was added and the cells were incubated for 2 h. [^3^H]-uridine (20 μCi/ml) was then added and the cells were incubated for a further 3 h., at which time the monolayers were washed 3 times with ice-cold PBS, lysed and RNA extracted with TRIzol reagent.

For measurement of viral RNA synthesis, the harvested RNAs were separated on a MOPS-Formaldehyde gel. The gel was fixed (15% methanol, 10% acetic acid and 75% dH2O) for 30 min followed by fluorography (Fluorographic reagent amplify, GE HEALTHCARE) for another 30 min. Gels were dried for 2 h. before exposure to autoradiographic film at −80 °C for 4 days.

For gradient analysis, equal volumes of harvested RNAs were loaded onto 14 ml 5-25% sucrose gradients in 100mM sodium acetate and 0.1% SDS followed by centrifugation at 150,000×g for 5 h. at room temperature. Gradients were fractionated into 350 μl fractions, and radioactivity of each fractions was determined by liquid scintillation counting.

### Analysis of CHIKV protein expression

C2C12 cells were electroporated with ICRES RNAs and incubated for 36 h. Cells were washed 3 times with PBS, lysed by resuspension in Glasgow lysis buffer (GLB) [1% Triton X-100, 120 mM KCl, 30mM NaCl, 5mM MgCl2, 10% glycerol (v/v), and 10 mM piperazine-N,N’-bis (2-ethanesulfonic acid) (PIPES)-NaOH, pH 7.2] supplemented with protease inhibitors and phosphatase inhibitors (Roche Diagnostics), and incubated on ice for 15 min. Following separation by SDS-PAGE, proteins were transferred to a polyvinylidene fluoride (PVDF) membrane and blocked in 50% (v/v) Odyssey blocking buffer (LiCor) in Tris-buffered saline (TBS) [50 mM Tris, 150 mM NaCl, pH 7.4]. The membrane was incubated with primary antibody in 25% (v/v) Odyssey blocking buffer overnight at 4°C, then incubated with fluorescently labelled anti-rabbit (800nm) secondary antibodies for 1 h at room temperature (RT) before imaging on a LiCor Odyssey Sa fluorescence imager.

### Expression and purification of AUD proteins

The wildtype and P247A/V248A AUD (nsP3 residues Ile141 to Gly374) were cloned into pET-28a-His-sumo for expression in *Escherichia coli* and subsequent analysis. His-sumo tagged AUD expression plasmids were transformed into Rosetta 2 and cultures were grown in Luria-Bertani (LB) medium supplemented with 50 μg/μl ampicillin and 1% (wt/vol) glucose. The cells were grown at 37°C to an optical density at 600 nm (OD600) of 0.5 to 0.7 and then induced with IPTG (isopropyl-D-thiogalactopyranoside) (0.5 mM) for 5 h at 18°C. The cells were harvested by centrifugation at 7,000 rpm for 10 min. Wildtype or mutant AUD protein was purified by sequential His-tag affinity purifications. Briefly, cell pellets were suspended in 20 ml AUD lysis buffer (100 mM Tris-HCl pH 7, 200 mM NaCl, 20 mM imidazole) supplemented with 2 μg/μl DNase and EDTA-free protease inhibitor cocktail tablets (Roche). The cell suspension was lysed by sonication on ice at an amplitude of 10 μm for six pulses of 20 s separated by 20 s and the extract clarified by centrifugation at 16000×g for 30 min at 4°C. The supernatant was filtered through a 0.45 μm filter and applied to a Ni^2^+ His-tag column for purification. Purified proteins were dialyzed to remove imidazole and sumo-protease was added to cleave the His-sumo tag. After dialysis, the proteins were applied again to the His-tag purification column, and the flow-throughs were collected as purified AUD proteins.

### Circular Dichroism (CD) Spectroscopy

Far-UV CD spectroscopy was performed on an APP Chirascan CD spectropolarimeter to obtain the secondary structure of AUDs. Spectra (190-260 nm) were recorded using 200 μl protein solution (at a concentration of 0.2 mg/ml) in a 1 mm path-length cuvette. Protein CD spectra deconvolution was analysed by DichroWeb.

### RNA filter binding assay

Radiolabelled RNA transcripts and AUD proteins were diluted in binding buffer (40 mM Tris-HCl [pH 7.5], 5 mM MgCl2, 10 mM DTT, 50 μg/ml bovine serum albumin, 10 μg/ml yeast tRNA [Ambion]) and pre-incubated separately for 10 min at 4°C. The binding reaction was initiated by mixing 1 nM radio-labelled RNA and AUD proteins (0 to 500 nM) in a 200 μl final volume at 4°C for 30 min. Membranes were pre-soaked in binding buffer supplemented with 5% (v/v) glycerol and assembled from bottom to top as follows in a slot-blot apparatus (Bio-Rad): filter paper, Hybond-N nylon (Amersham Biosciences) to bind free RNA molecules, and nitrocellulose (Schleicher & Schuell) to trap soluble protein-RNA complexes. After assembly, 200 μl of each binding reaction mixture was applied to each slot and filtered through the membranes. Each slot was washed with 0.5 ml of binding buffer and air dried, and quantification of radioactivity was performed using an image plate, BAS 1000 Bioimager (Fuji), and Aida Image Analyser v4.22 software. Fitting was performed using GraphPad Prism 5 software. In each case, the data were fitted to the hyperbolic equation R = Rmax x R/(Kd + [P]), where R is the percentage of bound RNA, Rmax is the maximal percentage of RNA competent for binding, [P] is the concentration of AUD, and Kd is the apparent dissociation constant.

### Precipitation of nsP3 and viral RNA

Co-precipitation experiments were performed in C2C12 cells electroporated with ICRES One-Strep-tag (OST) RNAs using Streptactin-agarose (Thermo Fisher Scientific), following the manufacturers protocol. Precipitated proteins were subjected to immunoblotting and co-precipitated RNAs were extracted by TRIzol and quantified by qRT-PCR.

### Distribution of nsP3, capsid protein and dsRNA in cells during CHIKV replication

Wildtype and nsP3-P247A/V248A CHIKV were introduced into ICRES-nsP3-ZsGreen-CHIKV where ZsGreen was fused into the hypervariable domain of nsP3. C2C12 cells were electroporated and harvested at defined times post electroporation, fixed with 4% paraformaldehyde (PFA), permeabilised by treatment with methanol, blocked with 2% BSA, and incubated with capsid protein antibody (gift from Andres Merits) or dsRNA antibody (J2 antibody, Scicons) at 4°C overnight, followed by secondary antibodies (Alexa Fluor 633 conjugated chicken anti-rabbit IgG and Alexa Fluor 594 conjugated donkey anti-mouse IgG) for 1h at room temperature. Distribution of nsP3, capsid protein and dsRNA were detected using a Zeiss LSM880 with Airyscan. Post-acquisition analysis was conducted using Zen software (Zen version 2015 black edition 2.3, Zeiss) or Fiji (v1.49) software [29].

### Co-localisation analysis

For co-localisation analysis, Manders’ overlap coefficient was calculated using Fuji ImageJ software with Just Another Co-localisation Plugin (JACoP) (National Institutes of Health) [30]. Coefficient M1 indicated here reports the fraction of the nsP3 signal that overlaps either the anti-dsRNA or anti-capsid signal. Coefficient values range from 0 to 1, corresponding to non-overlapping images and 100% co-localisation images, respectively. Co-localisation calculations were performed on >5 cells from at least two independent experiments.

### Statistical analysis

Statistical analysis was performed using unpaired two-tailed Student’s t tests, unequal variance to determine statistically significant differences from the results for the wild type (n≥3). Data in bar graphs are displayed as the means ± S.E.

## Supporting Information

**Supplementary Figure 1**. Alignment of AUD amino acid sequences (nsP3 residues 210-276) and ribbon structure of Sindbis virus AUD showing location of mutated residues (PDB ID code 4GUA).

**Supplementary Figure 2**: Sequence analysis of virus passage P0.

